# JEPEGMIX2: improved gene-level joint analysis of eQTLs in cosmopolitan cohorts

**DOI:** 10.1101/126300

**Authors:** Chris Chatzinakos, Donghyung Lee, Bradley T Webb, Vladimir I Vladimirov, Kenneth S Kendler, Silviu-Alin Bacanu

## Abstract

**Motivation:** To increase detection power, researchers use gene level analysis methods to aggregate weak marker signals. Due to gene expression controlling biological processes, researchers proposed aggregating signals for expression Quantitative Trait Loci (eQTL). Most gene-level eQTL methods make statistical inferences based on i) summary statistics from genome-wide association studies (GWAS) and ii) linkage disequilibrium (LD) patterns from a relevant reference panel. While most such tools assume homogeneous cohorts, our **G**ene-level **J**oint **A**nalysis of functional SNPs in **C**osmopolitan **C**ohorts (JEPEGMIX) method accommodates cosmopolitan cohorts by using heterogeneous panels. However, JEPGMIX relies on brain eQTLs from older gene expression studies and does not adjust for background enrichment in GWAS signals.

**Results:** We propose JEPEGMIX2, an extension of JEPEGMIX. When compared to JPEGMIX, it uses i) cis-eQTL SNPs from the latest expression studies and ii) brains specific (sub)tissues and tissues other than brain. JEPEGMIX2 also i) avoids accumulating averagely enriched polygenic information by adjusting for background enrichment and ii), to avoid an increase in false positive rates for studies with numerous highly enriched (above the background) genes, it outputs gene q-values based on Holm adjustment of p-values.

**Supplementary information:** Supplementary material is available at Bioinformatics online.

## 1 INTRODUCTION

Gene expression, is believed to have influenced human evolution and play a key role in diseases (Cookson, et al., 2009; Emilsson, et al., 2008; Kudaravalli, et al., 2009). Thus, it is critical for understanding diseases and developing treatments. The importance of gene expression was empirically supported by the enrichment of association signals in SNPs tagging gene expression (Ehret, et al., 2012; Fehrmann, et al., 2011; Fernandez, et al., 2012; Nica and Dermitzakis, 2008; Nicolae, et al., 2010), which are generally known as expression quantitative trait loci (eQTL).

Currently, the identification of complex disease susceptibility loci is performed via genome-wide association studies (GWAS). It involves scanning single nucleotide polymorphisms (SNPs) across the entire genome for genetic variations associated with a disease. Univariate analysis of GWAS is still the de facto tool for identifying trait/disease-associated genetic variants (Wellcome Trust Case Control, 2007). However, when analyzing more complex GWAS SNPs with weak or moderate effect sizes, the significant findings account only for a small fraction of the total phenotypic variation (Manolio, et al., 2009). Due to their small effect sizes, these variants are difficult to be detected in GWAS (Yang, et al., 2010). To increase the power of detection, researchers proposed analyzing genetic variants multivariately (Wang, et al., 2007).

One type of multivariate analyses is the transcriptome-wide association study (TWAS) which identifies significant expression-trait associations. Such methods, e.g. joint effect on phenotype of eQTL/functional SNPs associated with a gene (JEPEG) (Lee, et al., 2015), PredictXcan (Gamazon, et al., 2015), JEPEGMIX (Lee, et al., 2016) and TWAS (Gusev, et al., 2016) use eQTL to predict gene expression and/or infer which genes are associated with traits. However, unlike competing non-eQTL paradigms, e.g. LDscore/LDpred (Bulik-Sullivan, et al., 2015), current TWAS methods i) lack competitive adjustment for background enrichment (“average signal”) and ii) do not output q-values that control false positive rates when there is a substantial number of genes enriched (above background) in signals.

To address these shortcomings, we propose JEPEGMIX2, an extension of JEPEGMIX, which, in addition to the existing advantage of imputing eQTLs statistics and inferring gene-trait association in cosmopolitan cohorts, it also i) adjusts for background enrichment ii), offers the option to upweight rarer eQTLs and iii), to avoid false positive rate increase for high signal enrichment, it outputs Holm q-values.

## 2 METHOD

To avoid a mere accumulation of just averagely enriched polygenic information, we competitively adjust *χ*^2^ statistics for background enrichment (Text S1 Supplementary data (SD)). This is achieved by adjusting the statistic for average non-centrality. Such “centralized” JEPEGMIX statistic we simply denote as competitive (C), while the original statistic we deem as the non-competitive (NC) version. To facilitate user-specific input along with future extensions, the new annotation file now includes a R-like formula for the expression of each gene as a function of its eQTL genotypes (https://github.com/Chatzinakos/JEPEGMIX2). The annotation file includes cis-eQTL for all tissues available in PREDICTDB (http://predictdb.hakyimlab.org/). To avoid making inference about genes poorly predicted by SNPs, for the 44 available tissues we retain only genes for which the expression is predicted with *q*-value < 0.05 from its eQTLs. Additionally, given the increased deleteriousness of rarer mutations, we offer the possibility to upweight coefficient of rarer variants (Text S2 of SD for statistic computation) using a Madsen and Browning type approach (Madsen and Browning, 2009). For linkage disequilibrium (LD) estimates in cosmopolitan cohorts (needed for both imputation and statistical inference), we allow user to input the study cohort proportions of ethnicities from the reference panel. LD patterns of the study cohort are estimated as a weighted mixture (with the above weights) of the LD matrices for all ethnic groups in a reference panel (Text S3 of SD). LD patterns are subsequently used to i) accurately impute summary statistics of unmeasured eQTLs (Text S4 of SD) and ii), compute the variance of the SNP linear combinations used for gene level tests in each tissue (Text S3 of SD).

The current version uses the 1000 genome (1KG) Phase I release version 3 database as reference panel (Durbin, et al., 2010). It consists of 379 Europeans, 286 Asians, 246 Africans and 181 inhabitants of the Americas.

## 3 SIMULATIONS

To estimate the Type I error (false positive) rates of JEPEGMIX2, for five different cosmopolitan studies scenarios (Text S5 of SD), we simulated (under *H*_0_) 100 cosmopolitan cohorts of 10,000 subjects for Ilumina 1M autosomal SNPs using 1000 Genomes (1KG) haplotype patterns (Text S5, Table 1 of SD). The subject phenotypes were simulated independent of genotypes as a random Gaussian sample. SNP phenotype-genotype association summary statistics, were computed as a correlation test. Under *H*_0_, the resulting summary statistics are distributed as Gaussian. We obtained JEPEGMIX2 statistics for: i) competitive (C), non-competitive (NC) and ii) tests with rare (Madsen and Browning like) (R) and non-rare (NR) eQTL weights. To test the ability of methods to maintain false positive rates under background enrichment, we provide an enriched scenario (ES) where add a mean of one to the simulated “central” Z-score (CZ) to obtain “non-central” Z-score (NCZ) scenario. We also applied JEPEGMIX2 to 16 real summary datasets (Text S6, Table 2 of SD). To limit the increase in Type I error rates of JEPEGMIX2, due to certain genes being non-casual but just reasonably close to GWAS peaks, we deem as significantly associated only genes with Holm-adjusted *p*-value (*q*) < 0.05. Due to C4 explaining most of the signals in Major Histocompatibility (MHC) region (chr6: 25-33 Mb (McCarthy, et al., 2016), for schizophrenia (SCZ) we omit all non-C4 genes in this region.

## 4 RESULTS

JEPEGMIX2 with competitive (C) statistics, controls the false positive rates at or below nominal thresholds for both central (CZ) and non-central (NCZ) scenarios while the non-competitive (NC) has similar behavior only for the central case (when the GWAS statistics are not enriched) (Text S5, Figure S1-S5 in SD). Under the enriched scenario (NCZ) the non-competitive version of the test has much increased false positive rates.

Using the Holm p-value adjustment and both rare (R) and non-rare (NR) e QTL weights, for the real datasets significant gene signals were found in 9 traits, for which we present heatmaps (Text S6, FigureS6-S23in SD). Table1 shows the number of genes with, *q* – *value* < 0.05, in the real datasets (for the abbreviations see Table 2 in the SD).

**Table 1.** Number of Genes with *q* – *value* < 0.05 for the real data (see Table S2 for abbreviations).

| Traits | No Genes |
| --- | --- |
| SCZ | 68 |
| ALZ | 34 |
| AMD | 17 |
| BIP | 11 |
| HDL | 79 |
| LDL | 78 |
| T2D | 5 |
| TG | 48 |
| Smoking | 5 |

On a computational node with 4x Intel Xeon 6 core 2.67-GHz processor and 64GB of RAM, the single core schizophrenia JEPEGMIX2 run required slightly under 3.5 hours, including imputation of unmeasured eQTLs. For the remaining datasets the running time was below 3 hours.

## 5 CONCLUSIONS

We propose JEPEGMIX2, an updated software/method for testing the association between (cis-eQTL predicted) gene expression and trait. Unlike existing methods, even for highly enriched GWAS, JEPEGMIX2 competitive version fully controls the Type I error rates at or below nominal levels. To the desirable JEPEGMIX imputing/analyzing capabilities of cosmopolitan cohorts, we, thus, add a competitive version and extend the number of included i) eQTLs and ii) tissues. Unlike the previous version (and all competitors), it also accommodates up weighting of the rare variants and, for high enrichment (above background), it avoids the increased rate of false positives incurred by FDR p-value adjustment by employing a Holm adjustment. Being written in C++, JEPEGMIX2 has very low computational burden. Future versions of the software will be updated to use larger reference panels.

## REFERENCES

Bulik-Sullivan, B.K., et al. LD Score regression distinguishes confounding from polygenicity in genome-wide association studies. Nat. Genet 2015;47(3):291–295.

Cookson, W., et al. Mapping complex disease traits with global gene expression. Nat Rev Genet 2009;10(3):184–194.

Durbin, R.M., et al. A map of human genome variation from population-scale sequencing. Nature 2010;467(7319):1061–1073.

Ehret, G.B., et al. A multi-SNP locus-association method reveals a substantial fraction of the missing heritability. Am J Hum Genet 2012;91(5):863–871.

Emilsson, V., et al. Genetics of gene expression and its effect on disease. Nature 2008;452(7186):423–428.

Fehrmann, R.S., et al. Trans-eQTLs reveal that independent genetic variants associated with a complex phenotype converge on intermediate genes, with a major role for the HLA. PLoS Genet 2011;7(8):e1002197.

Fernandez, R.M., et al. Four new loci associations discovered by pathway-based and network analyses of the genome-wide variability profile of Hirschsprung’s disease. Orphanet J Rare Dis 2012;7:103.

Gamazon, E.R., et al. A gene-based association method for mapping traits using reference transcriptome data. Nat Genet 2015;47(9): 1091–1098.

Gusev, A., et al. Atlas of prostate cancer heritability in European and African-American men pinpoints tissue-specific regulation. Nat Commun 2016;7:10979.

Kudaravalli, S., et al. Gene expression levels are a target of recent natural selection in the human genome. Mol Biol Evol 2009;26:649–658.

Lee, D., et al. JEPEG: a summary statistics based tool for gene-level joint testing of functional variants. Bioinformatics 2015;31(8):1176–1182.

Lee, D., et al.JEPEGMIX: gene-level joint analysis of functional SNPs in cosmopolitan cohorts. Bioinformatics 2016;32(2):295–297.

Madsen, B.E. and Browning, S.R. A groupwise association test for rare mutations using a weighted sum statistic. PLoS. Genet 2009;5(2):e1000384.

Manolio, T.A., et al. Finding the missing heritability of complex diseases. Nature 2009;461(7265):747–753.

McCarthy, S., et al. A reference panel of 64,976 haplotypes for genotype imputation. Nat Genet 2016;48(10):1279–1283.

Nica, A.C. and Dermitzakis, E.T. Using gene expression to investigate the genetic basis of complex disorders. Hum Mol Genet 2008;17:R129–R134.

Nicolae, D.L., et al. Trait-associated SNPs are more likely to be eQTLs: annotation to enhance discovery from GWAS. PLoS Genet 2010;6(4):e1000888.

Wang, K., Li, M. and Bucan, M. Pathway-based approaches for analysis of genomewide association studies. Am J Hum. Genet 2007;81(6):1278–1283.

Wellcome Trust Case Control, C. Genome-wide association study of 14,000 cases of seven common diseases and 3,000 shared controls. Nature 2007;447(7145):661–678.

Yang, J., et al. Common SNPs explain a large proportion of the heritability for human height. Nat Genet 2010;42(7):565–569.

